# Optimization of High Molecular Weight DNA Extractions from Dried, Museum-Grade Insects Enables Long-Read Sequencing, Phylogenetics, and Methylation Profiling

**DOI:** 10.64898/2026.07.16.739004

**Authors:** Gabrielle A. Hartley, Richard J. Green, Nicole Pauloski, Nicole M. Tillquist, Phoebe Johnston, Sara Ord, Rachel J. O’Neill

**Author notes:** **Corresponding author:** Rachel J. O’Neill < >. denotes authors contributed equally (Gabrielle A. Hartley, Richard J. Green).

## Abstract

Developing an effective DNA extraction method that meets requirements for long-read sequencing of poorly preserved samples, such as museum specimens or ancient material, offers new opportunities for genomic analysis of endangered or extinct species for which samples are rare. However, these samples often yield degraded and highly fragmented DNA, rendering long-read sequencing infeasible for many specimens residing in museum collections. Herein, we demonstrate a protocol for successfully extracting DNA of sufficient quality for sequencing on the Oxford Nanopore Technologies long-read sequencing PromethION platform from a desiccated, museum-grade blue carpenter bee specimen (*Xylocopa caerulea*). We find the protocol is reproducible across specimens and yields high levels of long, endogenous *X. caerulea*-derived DNA, highlighting the utility of our method for enabling genomic studies of historical collections. From a single flow cell, we assembled the full-length mitochondrial genome and used this assembly to perform a phylogenetic analysis, accurately placing our *X. caerulea* specimen among related *Xylocopa* species, thus demonstrating the phylogenetic utility of long-read museomics. Using these long-read data, we analyzed native CpG methylation, finding endogenous methylation signals that correlate with genic and exonic sequences. This method expands the feasibility of genomic and epigenomic analyses from challenging samples, enhancing our ability to investigate the genomes of endangered and extinct species through archival resources.

## Introduction

Millions of preserved specimens are housed in natural history museums around the world, providing an abundant source of material for DNA sequencing and genomic analysis (Nachman et al. 2023; Lalueza-Fox 2022). However, the quality of DNA obtained from these specimens is affected by a multitude of sampling and storage conditions, including the original handling and collection of the specimen, tissue storage solution, exposure to UV light, temperature, pH, moisture, and humidity, often resulting in low molecular weight DNA (Zimmermann et al. 2008; Raxworthy and Smith 2021). DNA sequencing and analysis of specimens in museum collections are often relegated to short-read sequencing platforms due to sample conditions (Card et al. 2021; Holmquist et al. 2025); however, long-read sequencing platforms, such as Oxford Nanopore Technologies (ONT), offer significant advancements over short-read sequencing, such as improvements to genome assembly, increased resolution of structural variants and repetitive regions, complete mtDNA assemblies, and native detection of epigenetic modifications (Amarasinghe et al. 2020). As such, long-read sequencing of museum samples offers a compelling alternative to traditional short-read sequencing if challenges to DNA recovery can be surmounted.

In an effort to improve the DNA recovery from non-ideal specimens, we present a protocol for high-molecular weight (HMW) DNA extraction and long-read sequencing for desiccated and mounted samples of the blue carpenter bee, X*ylocopa caerulea*. Representing large carpenter bees, *Xylocopa* is a diverse genus containing roughly 400 species that vary widely in their coloration, distribution, and behavior (LaBerge 1965; Gerling et al. 1989; Leys et al. 2002). *X. caerulea* is a sexually dimorphic species found widely distributed throughout Asia, with females known for their distinct blue hairs of the thorax, abdomen, and head (Figure 1A), while males are largely black and uniform (Gu et al. 2023; Stavenga 2023). While recent studies have examined the gut microbiome (Gu et al. 2023) and coloration of *X. caerulea* (Stavenga 2023), little is known about the social behavior, genetics, or epigenetics of this species and there is no genomic data publicly available for *X caerulea*.

**Figure 1:**
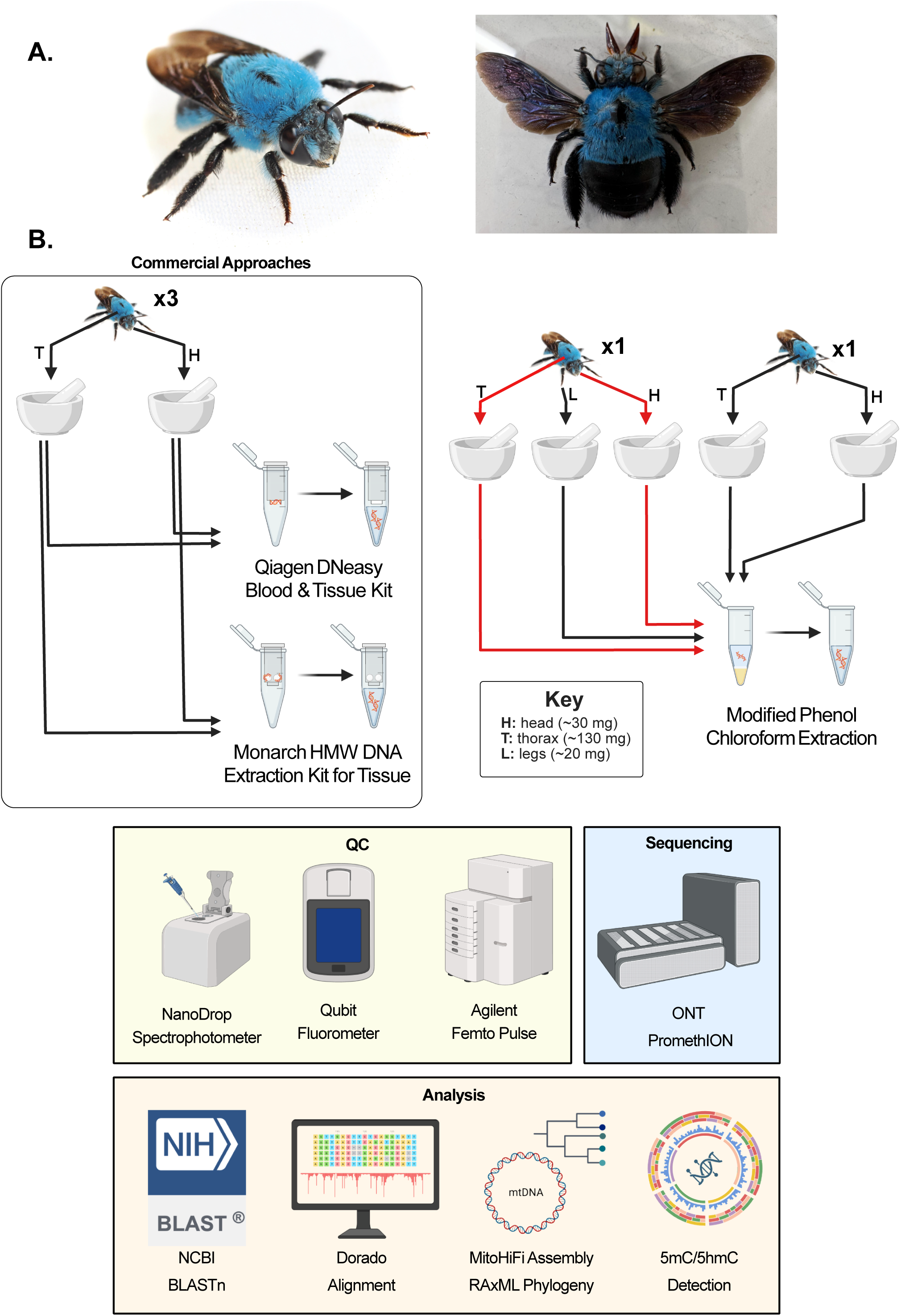
Experimental workflow for long-read ONT sequencing and analysis of a museum-grade *X. caerulea* specimen. **(A)** A live *Xylocopa caerulea* specimen (left) and an example of a pinned and mounted *X. caerulea* specimen obtained for sequencing in this study (right). **(B)** Overview of the sample preparation, extraction, quality control, sequencing, and downstream analysis methods used in this study. High molecular weight (HMW) DNA was isolated independently from the head (H; ∼30 mg), thorax (T; ∼130 mg), and legs (L; ∼20 mg) of *X. caerulea* specimens using either commercial DNA extraction kits (Monarch HMW DNA Extraction Kit for Tissue and Qiagen DNeasy Blood & Tissue Kit) or a modified phenol-choloform extraction protocol. DNA quantity and quality were assessed using the NanoDrop spectrophotometer, Qubit fluorometer, and Agilent Femto Pulse system (yellow). Based on DNA quality, quality, and length, the thorax (T) and head (H) samples extracted using the modified phenol chloroform extraction (red arrows) were selected for sequencing on the Oxford Nanopore Technologies PromethION platform (blue). Following sequencing, we used a BLASTn search and whole genome alignment to classify read origin, RAxML to infer phylogenetic placement, and base modification analyses to detect 5-methylcytosine (5mC) and 5-hydroxymethylcytosine (5hmC) modifications (orange).

Using our newly developed protocol, we confirmed the presence of HMW DNA fragments and successfully generated long-read data on the ONT platform, demonstrating the recovery of long, endogenous *X. caerulea* DNA despite non-ideal specimen storage conditions (Figure 1B). We used these sequencing results to create a high-quality mitochondrial assembly and performed a phylogenetic analysis with the *COX1* gene, correctly placing this specimen among closely related *Xylocopa* species. Lastly, we leveraged this long-read sequencing data to detect native CpG methylation in *X. caerulea*, highlighting the capacity of long-read sequencing to simultaneously capture genetic and epigenetic data from historical specimens. Collectively, these protocols expand the range of specimens compatible with long-read sequencing platforms, enabling genetic and epigenetic analyses of rare, degraded, or otherwise challenging samples preserved in natural history collections.

## Materials and Methods

### Sample Collection

A total of five *X. caerulea* specimens were initially purchased from the online retailer Amazon. They arrived desiccated, having previously been pinned and mounted for display. Prior to DNA extraction, the contents of the abdomen and the wings were removed to minimize recovery of endosymbiont and gut-derived DNA. Samples were ground in a mortar and pestle in liquid nitrogen. Three methods of DNA extraction were chosen: Qiagen DNeasy column extraction, Monarch HMW DNA extraction, and a modified version of (Green and Sambrook 2012) phenol-chloroform extraction (described below and in Supplementary Table 1).

### DNA Extractions

#### Extraction Method 1: Qiagen DNeasy Blood and Tissue Kit Extraction

DNA extraction was performed using the Qiagen DNeasy Blood and Tissue Kit (Qiagen #69504) according to the manufacturer’s instructions. A total of two DNA extractions were performed with this method using pooled thorax and pooled head tissue from three specimens each (Supplementary Table 2). The final DNA was eluted in 200μl of Buffer AE.

#### Extraction Method 2: New England BioLabs Monarch HMW DNA Extraction

DNA extraction was performed using the New England BioLabs Monarch HMW DNA Extraction Kit (NEB #T3060) protocol for tissue according to the manufacturer’s instructions. A total of two DNA extractions were performed with this method using pooled thorax and pooled head tissue from three specimens each (Supplementary Table 2). During lysis incubation, the samples were agitated at 650 rpm in the thermal mixer, agitating with a micro-pestle every 15 minutes.

#### Extraction Method 3: Phenol-Chloroform Extraction

DNA was extracted following the Green and Sambrook’s phenol-chloroform extraction protocol with modifications (Supplementary Table 1). A total of five DNA extractions were performed with this method (Supplementary Table 2). First, three individual extractions were completed with thorax, head, and leg tissue from a single specimen; subsequently, two individual extractions were completed with thorax and head tissue to replicate the results. To precipitate the DNA, one-half volume of 3M ammonium acetate was gently added to the aqueous solution and mixed to combine, followed by the addition of 2.5 volumes of 100% cold ethanol. Samples were mixed gently by inverting into solution and incubated at 4°C overnight. The precipitant DNA was centrifuged at 8,000 x g for 30 minutes. Following centrifugation, all but 6 mL was removed, gently mixed and transferred to a 2ml tube in 2ml aliquots and centrifuged at 10,000 x g for 20 minutes. The supernatant was removed, and this process was repeated for the remaining solution. The DNA pellet was then washed twice with 1ml of 70% ethanol and centrifuged at 10,000 x g. The ethanol was removed and the pellet was air-dried for approximately 10 minutes. The DNA was eluted in 100μl of TE buffer and left at room temperature (RT) overnight.

### DNA Quantification and Size Selection

DNA concentrations were measured using the dsDNA high sensitivity kit (cat# Q33230) on an Invitrogen Qubit 4 Fluorometer per manufacturer’s instructions. The dsDNA purity was measured using a Thermo Scientific Nanodrop One Spectrophotometer. DNA fragment sizes were measured using both the Agilent TapeStation and Femto Pulse Genomic DNA 165 kit (cat# FP-1002-0275).

Prior to library preparation, the purified libraries from head and thorax phenol-chloroform extraction(R5-R6, Supplementary Table 2) were pooled and size selected using Agencourt Ampure XP beads to enrich for DNA fragments above 1.5-2 kb. Beads were used at a ratio of 35μl beads to 50μl resuspended DNA and gently flicked to mix.

### ONT Library Preparation

Library preparation was performed using ONT’s DNA V14 (SQK-LSK114) kit with the following modifications. During *DNA repair and end prep*, the optional DNA Control Sample was not used. The reaction was incubated at 20°C for 20 minutes and 65°C for 10 minutes. After the addition of Ampure XP beads, the sample was incubated on a Hula Mixer for 10 minutes at room temperature (RT). After the ethanol washes, the sample was resuspended in 61μl of Nuclease-free water for 5 minutes at 37°C. For *Adapter ligation and clean-up*, we used the Short Fragment Buffer (SFB) and incubated the reaction for 20 minutes at room temperature. We added 100μl of resuspended AMPure XP beads to the reaction and incubated the sample on a Hula Mixer for 10 minutes. After the SFB wash and supernatant removal, the pellet was resuspended in 30μl of Elution Buffer (EB) and incubated for 10 minutes at 37°C. The beads were pelleted and 30μl of elute was retained, quantified and loaded onto a PromethION R10.4.1 flow cell per the standard LSK114 protocol (Supplementary Table 3). The flow cell was loaded 3 times across a 90-hour run period.

### DNA Content Analysis

The data was basecalled using the Dorado basecaller (v0.8.20) (*Dorado: Oxford Nanopore’s Basecaller*, n.d.) with a minimum Q score of 10 and the sup,5mCG_5hmCG model. Basecalled data was analyzed using NanoPlot (v1.34.1) (De Coster and Rademakers 2023) to determine QC metrics. To identify the origin of the reads (ie., endogenous *X. caerulea* vs. exogenous contamination), we conducted a BLASTn (v2.7.1) (Altschul et al. 1990) search of the reads using NCBI’s Basic Local Alignment Search Tool (BLAST) with the full NCBI non-redundant nucleotide database. The query coverage per high-scoring segment pair (qcov_hsp_perc) was set to 15% to minimize spurious short matches. Each sequence read was compared to the total length of alignments, average Q score, and top scoring BLAST match (by E value) to obtain accurate classifications. Reads matching the BLAST classification “bees,” “ants,” “wasps, ants, and bees,” and “hymenopterans” were considered a conservative estimate to identify endogenous, *Xylocopa*-derived reads (Supplementary Table 5, Supplementary Table 6). NanoPlot (v1.44.0) (De Coster and Rademakers 2023) and Seqkit (v2.10.044.0) (Shen et al. 2024) were used to compare read length and quality of endogenous reads.

### Mitochondrial Analysis

To assemble the mitochondria, we filtered ONT reads by length using Seqkit (v2.10.044.0) (Shen et al. 2024), retaining reads between 2,000 bp and 18,000 bp to reduce the likelihood of nuclear mtDNAs (NUMTs) in the analysis. Reads were assembled and annotated using MitoHiFi (v3.2.1) (Uliano-Silva et al. 2023; Allio et al. 2020) with the complete mitochondria of *Ceratina smaragdula* (NC_064404.1) as a reference, the closest phylogenetic relative available on NCBI.

### Alignment and Phylogenetic Analysis

Reads were aligned to four publicly available reference genomes for *Xylocopa* species (Supplementary Table 4) using the Dorado aligner (v0.8.20) (*Dorado: Oxford Nanopore’s Basecaller*, n.d.) and each alignment was indexed using Samtools (v1.20) (Li et al. 2009). Because our sequencing data aligned best to the *X. dejeanii* genome assembly (GCA_049004755.1) (Zhang et al. 2025), this reference was selected for subsequent analyses.

We collected complete and partial *COX1* gene sequences for 28 related species available on NCBI (Supplementary Table 7) and performed a multiple sequence alignment using the global alignment with free end gaps setting in the Geneious Prime suite (Kearse et al. 2012). Based on this alignment, the maximum sequence portion shared by all species was retained and used in phylogenetic analysis were constructed with RAxML-NG (v1.2.2) (Stamatakis 2014) with 1,000 bootstraps using the model GTR+I+G.

### Methylation Analysis

To analyze methylation, genomic scaffold names were standardized using *cut,* grep, and *awk* commands to filter the *X. dejeanii* genome assembly (GCA_049004755.1) (Zhang et al. 2025) to remove unplaced scaffolds. DNA modification calls (5mC and 5hmC) were generated using modkit (v0.2.5) pileup (*Modkit: A Bioinformatics Tool for Working with Modified Bases*, n.d.) in a CpG context. To facilitate genomic visualization, we separated modification calls by type (m vs. h) and converted into BedGraph format, representing the percentage of methylated or hydroxymethylated cytosines at each site. We analyzed broad-scale methylation patterns by binning the genome into fixed 100kb windows using bedtools (v2.31.1) makewindows (Quinlan and Hall 2010). Gene density, 5mC, and 5hmC methylation for each 100kb window was calculated using bedtools (v2.31.1) intersect (Quinlan and Hall 2010) and the resulting bedgraph tracks were loaded into RStudio (v4.3.1). To visualize the global methylome landscape, a circos plot was generated using the circlize (v0.4.17) (Gu et al. 2014) package. To ensure visibility across varying data ranges, tracks were auto-scaled to their respective maximum with a 10% headroom. The distribution of methylation marks across specific genomic features (5’ UTR, Exons, Introns, and 3’ UTR) was analyzed and visualized using ggplot2 (v4.0.1) (Wickham 2016). For high-resolution gene-body analysis, we used methylartist (v1.2.7) segmeth (Cheetham et al. 2022) to calculate the mean 5mC and 5hmC percentages specifically within the boundaries of each annotated gene. Locus-specific plots for top-ranking methylated genes were generated using methylartist (v1.2.7) locus (Cheetham et al. 2022), utilizing a smooth window size of 15. Functional identification of top methylated hits was performed by extracting gene sequences with samtools (v1.21) faidx (Li et al. 2009) and conducting homology searches via BLASTx (NCBI) (Altschul et al. 1990).

## Results

### DNA Extractions

To obtain DNA from *X. caerulea* specimens purchased at an online retailer, we first sought to determine if two commercially available kits for DNA extraction would be suitable for obtaining HMW DNA (Figure 1B, 1C). To that end, we independently pooled head and thorax tissue from three specimens and performed DNA extractions on each using the Qiagen DNeasy Blood and Tissue Kit and the Monarch HMW DNA Extraction Kit. Using the Qiagen DNeasy Blood and Tissue Kit, we obtained 1,164 and 1,084 ng of DNA from the pooled head and thorax tissue, respectively, providing enough DNA recovery for ONT sequencing (R1 and R2, Supplementary Table 2). However, when analyzed for fragment length, the DNA was suboptimal for long-read sequencing, with shorter primary fragment sizes and more contaminants than subsequent extraction methods, resulting in lower A260/230 and A260/280 ratios (Supplementary Figure 1, Supplementary Table 2). Using the Monarch HMW DNA Extraction kit, we obtained non-quantifiable DNA yield, rendering the method inadequate for long-read sequencing without amplification (R3 and R4, Supplementary Table 2). Therefore, we proceeded with a modified phenol-chloroform extraction to improve DNA yield and fragment size.

In total, we performed five phenol-cholorform DNA extractions utilizing various conditions. First, we performed extractions on three tissue types from a single specimen, including head, thorax, and leg tissue, yielding 3,680 ng, 16,500 ng, and 51.6 ng of DNA, respectively (R5-R7, Supplementary Table 2). Subsequently, extractions using head and thorax tissue were replicated to address potential variability in sample condition among specimens (R8 and R9, Supplementary Table 2), yielding 466 ng and 570 ng of DNA in head and thorax extractions, respectively.

Across all extractions, the modified phenol-chloroform extraction on multi-specimen thorax tissues provided the best fragment lengths, with average DNA size peak length up to 42,807 bp (Supplementary Table 2). In addition to increased fragment length, extractions from thorax tissue yielded more optimal A260/230 and A260/280 ratios (Supplementary Table 2). While subsequent repeated extractions yielded optimal A260/230 and A260/280 ratios and larger fragment lengths sufficient for ONT sequencing, they resulted in less total DNA than prior extractions, suggesting variability among specimens (Supplementary Table 2). As such, the extractions for thorax and head tissue (R5-R6, Supplementary Table 2) were selected as the best samples when comparing contamination, quantity, and fragment length. Thus, these two extractions (R5-R6, Supplementary Table 2) were pooled, size selected to remove DNA fragments smaller than 2kb, and prepared for sequencing.

#### Oxford Nanopore Technologies (ONT) Sequencing

Following library preparation, the sample was sequenced on an ONT PromethION R10.4.1 flow cell, generating approximately 11.17 million reads passing quality thresholds >Q10 and totaling ∼5.82 Gb, or roughly ∼30X coverage when compared to the closely related *X. dejeanii* genome (ASM4900475v1, 194.5 Mb) (Supplementary Table 3) (Zhang et al. 2025). Passed reads had a median quality score of 13.9 and an N50 of 546 basepairs (bp) with a standard deviation of 582 bp. To further assess the quality of our sequencing data, we aligned our sequencing reads to four publicly available reference genomes within the *Xylocopa* phylogeny (Supplementary Table 4). We found that our data had the highest alignment rate to the reference assembly for *X. dejeanii*, with an alignment rate of 11.33% (8.6%-11.33%) (Supplementary Table 4). When comparing genome-wide read distribution of our *X. caerulea* specimen to the genome of *X. dejeanii*, we found relatively uniform coverage across the *X. dejeanii* genome with the exception of a few chromosomes (Figure 2A); however, these *X. dejeanii* chromosomes with lower coverage are significantly shorter than the anticipated chromosome lengths, indicating potential misassemblies in the *X. dejeanii* genome that may affect read coverage in these regions. As such, this genome was selected for subsequent analyses, and mapped reads were extracted from the alignment.

**Figure 2:**
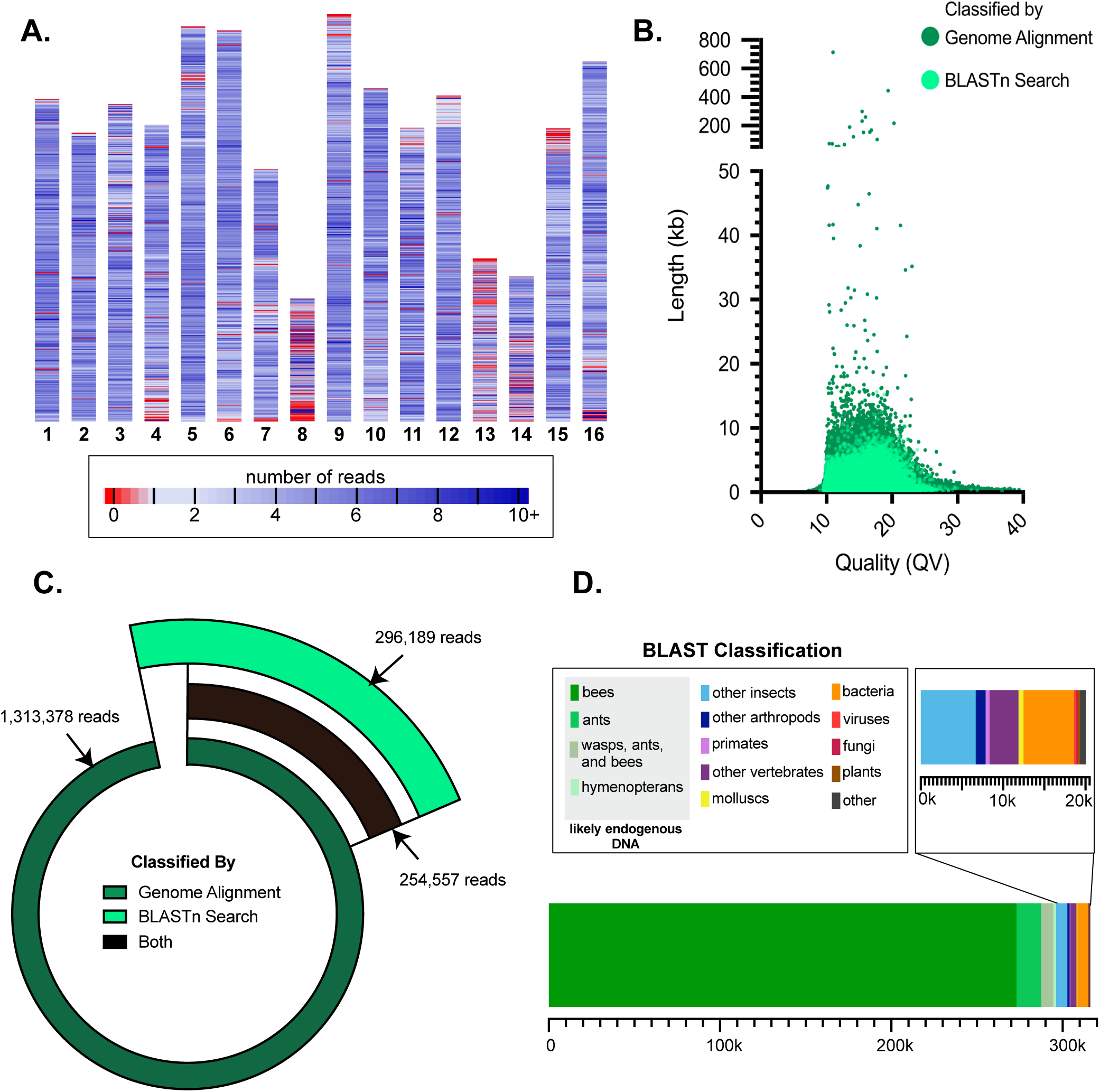
Classification and alignment of long-read ONT sequencing reads from an *X. caerulea* specimen. **(A)** Distribution of read depth across the *X. dejeanii* genome (GCA_049004755.1), showing generally even coverage across the genome. Across the chromosomes, the average read depth is indicated from low (1X depth, light blue) to high (10X+ read depth, dark blue). Regions with no coverage (red) are more frequent on misassembled chromosomes (as indicated by chromosome lengths that are shorter than expected). **(B)** Read length (kilobases, kb; y-axis) and quality (QV; x-axis) distribution for ONT-sequencing reads classified as endogenous via genome alignment to the *X. dejeanii* genome (dark green) and BLASTn search (light green). **(C)** Distribution of reads classified via genome alignment to the *X. dejeanii* genome (dark green), BLASTn search (light green), and both methods (black). 14% of reads were classified as endogenously derived via BLASTn search but did not align to the *X. dejeanii* genome, representing potential species-specific variation. **(D)** A summary of BLASTn taxonomic read classifications by read count with bar lengths indicating the number of reads assigned to taxonomic groups. *X. caerulea*-derived reads constitute the largest proportion of classified reads (green, 94%), herein categorized as anything classified with the taxonomic label “bees,” “ants,” “wasps, ants, and bees,” and “hymenopterans.” An insert panel displays exogenous read classifications, a minor fraction of all classified reads.

In total, roughly 1.3 million reads aligned to the *X. dejeanii* assembly with an N50 of 1,148 bp and a standard deviation of 1,230.8 bp, more than double the N50 of the overall sequencing run and representing a significant increase in read length when compared to traditional short-read approaches (Figure 2B). The average Q score of mapped reads was 14.5 and the longest mapped read was 712,506 bp. To identify *Xylocopa*-derived DNA in our sequencing data even more conservatively, we performed a BLASTn search of all reads produced to classify reads more broadly across all taxa. In total, 316,225 unique reads produced high quality alignments in a BLASTn search belonging to 258 unique taxonomic identifiers (taxids). Of these, 93.7% (296,189 reads) produced best hit alignments to BLAST classification groups “bees”, “ants,” “wasps, ants, and bees,” and “hymenopterans” (Figure 2C, 2D), which are herein classified as endogenous due to their phylogenetic relation to *Xylocopa* species. Reads classified as endogenous via BLASTn search had a read length N50 of 906 bp with a standard deviation of 637.8 bp (Figure 2B) and an average Q score of 14.1 (Figure 2B). While ∼86% of BLAST-classified reads were concordantly identified as endogenous in both classification strategies, an additional 41,632 reads were classified as endogenous via BLAST search only, potentially representing species-specific variation between *X. caerulea* and *X. dejeanii* (Figure 2C). Combined, the results of our genome alignment and BLASTn search indicate the presence of high-quality, endogenous DNA sequences from the museum-grade *X. caerulea* specimen.

#### Mitochondrial Assembly and Phylogeny

Mitochondrial analysis is commonly employed to provide genetic and phylogenetic insights from museum specimens by leveraging high copy mitochondrial DNA fragments. To that end, we sought to examine the feasibility of assembling a mitochondrial sequence from our *X. caerulea* ONT sequencing data. To reduce the likelihood of including NUMTs in our assembly while still utilizing the mappability benefits of long-reads, we filtered our ONT read set to retain fragments between 2 kilobases (kb) and 18 kb and generated a mitochondrial assembly using MitoHiFi (Uliano-Silva et al. 2023). The resulting mtDNA assembly is 17,451 bp and contains 37 genes with no frameshifts (Figure 3A), indicating successful recovery of mitochondrial sequences. To assess the accuracy of our assembled mitochondrial genome and to demonstrate the ability to perform phylogenetic analyses from long-reads derived from museum-grade specimens, we isolated the *COX1* gene from our assembly and obtained 28 full length and partial *COX1* gene sequences for bees available on NCBI (Supplementary Table 7). Using these sequences, we performed a multiple sequence alignment and generated a phylogenetic tree using RAxML (Stamatakis 2014) with 10,000 bootstraps (Figure 3B). We find that our assembled *X. caerulea COX1* sequence was placed amongst other *Xylocopa*-derived gene sequences, matching expected phylogenetic predictions for this species.

**Figure 3:**
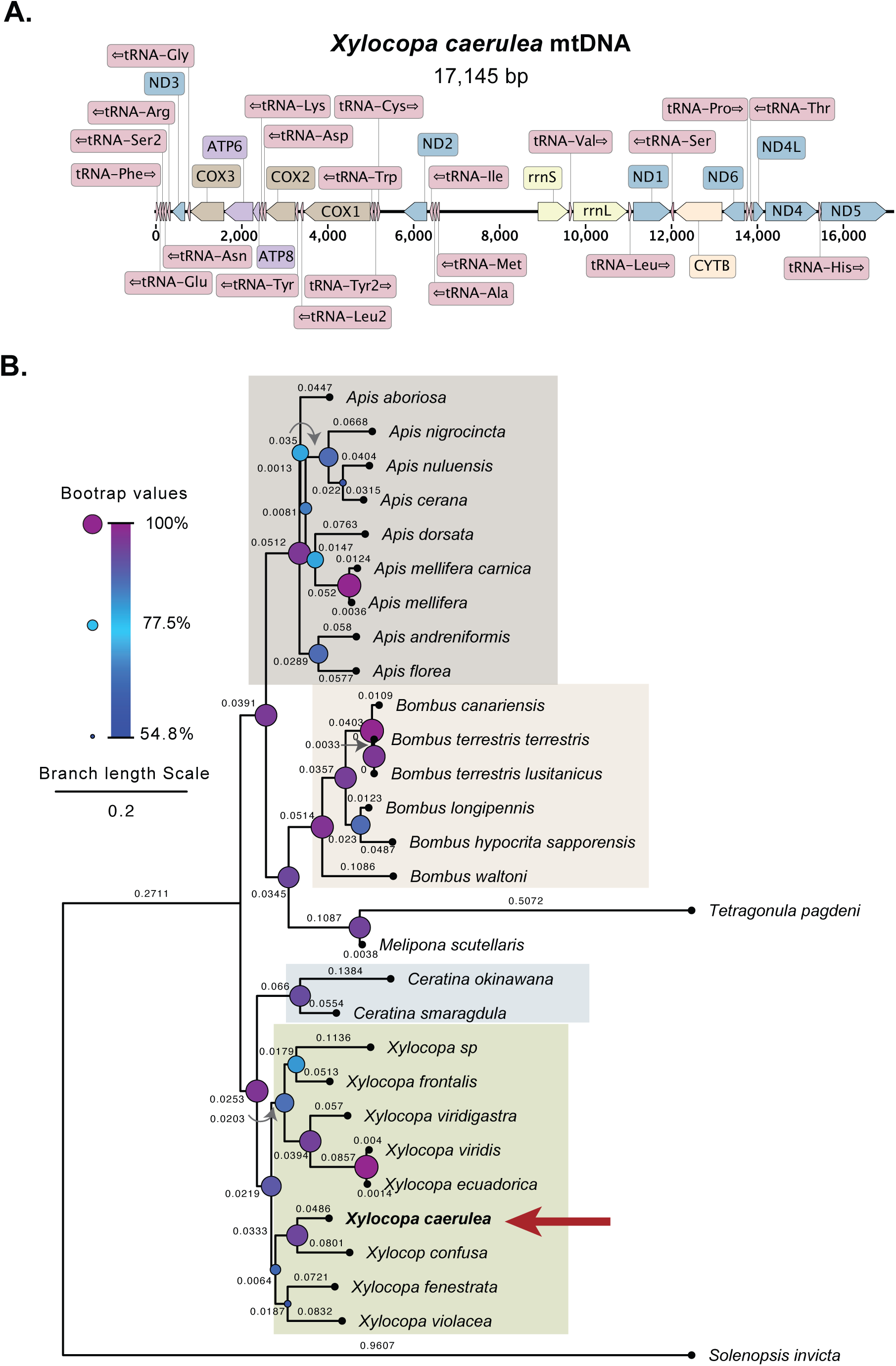
Mitochondrial genome organization and phylogenetic analysis of *X. caerulea*. **(A)** A linearized depiction of the 17,145 basepair (bp) mitochondrial DNA assembled from *X. caerulea* ONT reads. 37 genes were identified, including 13 protein coding genes (NADH dehydrogenase subunits, blue; cytochrome C subunits, brown; ATP synthase subunits, purple; and Cytochrome C, orange), 22 transfer RNAs (tRNAs, red), and 2 ribosomal RNAs (rRNAs, yellow). The assembly generated in this study provided a complete mitochondrial genome for *X. caerulea* suitable for subsequent phylogenetic analysis. **(B)** A maximum likelihood phylogeny inferred from the *X. caerulea COX1* sequence assembled herein and 28 publicly available *COX1* sequences (Supplementary Table 7), including 27 sequences from across the Apidae family and one sequence for *S. invicta*, the red imported fire ant. Numbers at the internal nodes indicate the branch lengths. Bootstrap values are represented by circles on the nodes ranging from low (blue, small, 54.8% minimum) to high (purple, large, 100% maximum). Boxes indicate the *Apis* (gray), *Bombus* (orange), *Ceratina* (blue), and *Xylocopa* (green) genera, with the *X. caerulea* specimen sequenced in this study correctly placed amongst the *Xylocopa* genera (red arrow).

#### Methylation

Epigenetic marks like CpG methylation play a crucial role in the maintenance of genome function and stability (Miller et al. 1974; Putiri and Robertson 2011). While resolving DNA methylation patterns could inform key biological processes across non-model species, obtaining epigenetic data from museum-quality specimens remains a challenge due to sample conditions, DNA degradation, and material availability. In bees, methylation patterns define the regulation of individual-specific traits, such as phenotypic plasticity and complex social behaviors displayed by members of the Apoidea superfamily (Lyko et al. 2010; Yagound et al. 2020; Rahman and Lozier 2023; Wang et al. 2020). Therefore, we decided to take advantage of the real-time epigenetic modification capture available using ONT sequencing to assess CpG methylation in *X. caerulea*. Using reads mapped to the closely related *X. dejeanii* reference assembly (GCA_049004755.1) (Zhang et al. 2025), we visualized the methylome of *X. caerulea* across the 16 chromosome scaffolds in 100 kb windows, finding that both 5mC and 5hmC modifications are broadly distributed across the genome (Figure 4A). While 5mC levels (blue, maximum enrichment ∼30%) were generally higher across windows than 5hmC (orange, maximum enrichment ∼1.2%), both modifications showed increased methylation corresponding with gene-dense regions (green) matching expectations for endogenous CpG methylation patterns in bees (Rahman and Lozier 2023) (Figure 4A, Supplementary Table 8).

**Figure 4:**
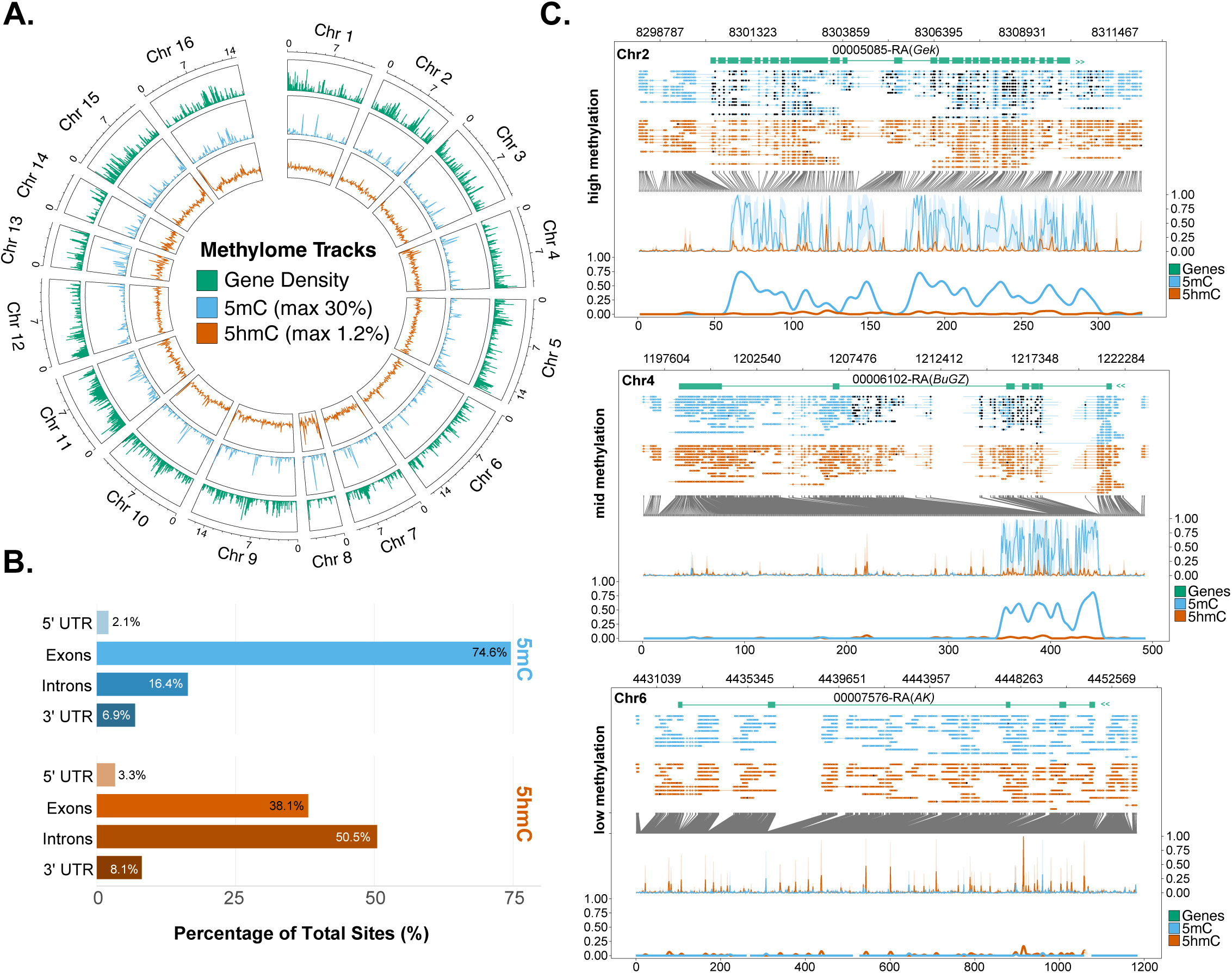
Epigenetic landscape and genomic distribution of 5mC and 5hmC DNA methylation modifications derived from a museum-grade *X. caerulea* specimen. **(A)** A circos plot illustrates the chromosome-wide distribution of gene density and methylation levels called by aligning *X. caerulea* reads to the *X. dejeanii* (GCA_049004755.1) genome. The plot depicts the gene density (green, outer track), 5mC levels (blue, middle track; max 30%), and 5hmC levels (orange, inner track; max 1.2%), showing that increases in gene density coincide with increases in methylation signatures. **(B)** Percentage of total modified 5mC (blue) and 5hmC (orange) sites across specific transcript features, including the 5’ UTR, exons, introns, and 3’ UTR. An increase in 5mC modifications is observed in exons, matching expectations for such epigenetic marks. Percentages are normalized to equal 100% per modification. **(C)** Representative single-locus genomic tracks illustrating methylation patterns across genes with high methylation (*Gek*, top), mid methylation (*BuGZ*, middle), and low methylation (*AK*, bottom). Tracks depict genes present on the loci (green), per strand 5mC (blue) and 5hmC (orange) modifications, and aggregate methylation frequency plots at the base of each panel. Variable patterns of epigenetic modifications observed across genes coincident with the presence of methylation marks consistent across multiple reads confirm the presence of endogenous, *X. caerulea*-derived methylation signatures.

Analysis of the distribution of methylation marks across specific genomic features (5’UTR, introns, exons, 3’UTR) revealed a distinct preference for gene bodies (Figure 4B). Among genomic features, the majority of 5mC sites (74.6%) were localized within exons, while 16.4% were found in introns; in contrast, 5hmC was more frequently associated with non-coding regions within the gene, with half of all 5hmC sites (50.5%) residing in introns compared to 38.1% in exons (Figure 4B). Consistent with literature reporting sparse CpG methylation within promoters complementing higher methylation in gene bodies among insects (Jeong et al. 2018; Lewis et al. 2020; Yagound et al. 2020), we found that both the 5’ and 3’ untranslated regions (UTRs) exhibited relatively low levels of modification compared with exons (Figure 4B). Despite this, CpG methylation was consistently detectable in 5’ and 3’ UTRs with three times as much 5mC enrichment in the 3’ UTR (6.9%) as the 5’ UTR (2.1%). A similar trend was observed for 5hmC (8.1% in 3’ UTR vs 3.3% in 5’ UTR), suggesting that epigenetic marking increases toward the distal end of transcriptional units.

To validate the reliability of methylation calls, we examined representative loci across three distinct levels of 5mC enrichment levels across the gene (high, mid, low 5mC modifications; Figure 4C, Supplementary Table 9). *Gek*, a protein kinase associated with actin polymerization (“Gek Genghis Khan [Drosophila Melanogaster (fruit Fly)] - Gene - NCBI,” n.d.), displayed generally even 5mC modifications across the gene body, with 5mC modifications consistently detected in most reads mapping to specific CpG sites. *BuGZ*, a gene critical for facilitating mitotic spindle assembly and microtubule binding (“BuGZ Bub3 Interacting GLEBS and Zinc Finger Domain Protein [Drosophila Melanogaster (fruit Fly)] - Gene - NCBI,” n.d.), similarly displayed 5mC modifications consistently detected across reads, but modifications were localized to the 5’ side of the gene body. In contrast to these loci, *Argk*, a gene related to arginine kinase activity (“Argk1 Arginine Kinase 1 [Drosophila Melanogaster (fruit Fly)] - Gene - NCBI,” n.d.), displayed low, sporadic 5mC levels. These differing epigenetic profiles demonstrate that our extraction and sequencing protocols effectively captured structured biological signals rather than stochastic noise.

## Discussion

Despite the availability of millions of specimens existing in worldwide sample collections, the recovery of high molecular weight DNA fragments from historical, museum, and ancient samples remains a significant challenge to the field of museomics. Such samples are prone to DNA degradation and contamination due to age and storage conditions and are therefore typically relegated to traditional short-read sequencing methods rather than emerging long-read sequencing methodologies commonly employed in modern genetic, genomic, and epigenetic studies (Zimmermann et al. 2008; Raxworthy and Smith 2021). Moreover, since DNA methylation analysis involves a specific preparation for short-read sequencing platforms (e.g. Illumina) independent of whole genome shotgun sequencing, the epigenetic landscape of samples residing in museum and natural history collections have remained largely unknown. Addressing these technical limitations is particularly essential in expanding genomic and epigenetic resources for species of ecological, agricultural, or conservation concern, which may reduce the ability for fresh sampling.

To demonstrate this utility, we obtained HMW DNA of sufficient length and quantity to successfully perform ONT long-read sequencing on dried and mounted specimens of *Xylocopa caerulea*, the blue carpenter bee, using our modified extraction protocol. From a single flow cell of data, we successfully performed comparative genomic analyses, including mitochondrial assembly and annotation, phylogenetics, and methylation profiling, thus allowing us to simultaneously place specimens in a phylogeny while capturing enough genome coverage to study gene and genome structure in the context of DNA methylation. Carpenter bees of the family *Xylocopa* have unique utility to agriculture given the ease of solitary bee population management and their increased pollination efficiency across a range of crop and non-crop plants (Eeraerts et al. 2020; Mensah and Kudom 2011; Junqueira et al. 2012). Recent advancements in crop productivity have come at a great cost to pollinator diversity, which in turn negatively impacts the holistic maintenance of entire ecosystems that depend on a wide variety of pollinator species, including the more elusive native species (Winfree et al. 2018; Deguines et al. 2014). A lack of genomic data for native pollinator species like *X. caerulea* thus limits the ability of the scientific community to confront the collapse of pollinator populations, and the accessibility of such data is further narrowed when many of the available specimens are stored in non-ideal conditions, such as dried and mounted.

Our ability to harness long-read whole-genome sequencing data, including native methylation profiling, from a dried and mounted sample provides a powerful means to address a diverse set of research aims, from methylation-based gene regulation among social and solitary behaviors in bees to the genetic basis of declining populations within pollinators to develop epigenetic-aware conservation strategies (Seebacher and Krause 2019; Rey et al. 2020). As a result, collections once considered suboptimal for such studies can now serve as new resources to investigate genetic diversity and genome regulation. Broadly, the rich reservoir of epigenetic and genetic information accessible using long-read sequencing data substantially enhances the future utility of dried, desiccated, or museum specimens previously relegated to short-read sequencing technologies.

## Acknowledgements

Within the Institute for Systems Genomics, the Center for Genome Innovation assisted with ONT PromethION work and the Computational Biology Core provided High Performance Computing (HPC) support. We gratefully acknowledge Timothy Earley for providing useful discussions that improved this work. The *Xylocopa caerulea*, blue carpenter bee image was provided by Dan Brown with Wild Discovery, www.wild-discovery.com.

## Data Accessibility Statement

Data associated with this manuscript can be accessed under BioProject PRJNA1466300 and the BioSample SAMN60045192. Long-read ONT sequencing can be accessed under SRA: SRR38613888. The mitochondrial assembly for *X. caerulea* can be accessed in GenBank using the accession PZ572862.

## Benefits Sharing Statement

Benefits Generated: This study established a reproducible method for recovering high-quality, long-read sequencing data from a dried museum specimen, expanding the genomic utility of historical collections. By enabling mitochondrial assembly, phylogenetic inference, and native methylation detection, this approach increases access to genomic and epigenetic data for rare, endangered, or extinct species for which samples cannot be readily obtained. Broadly, this work reflects the partnership between the UConn Institute for Systems Genomics and the Colossal Foundation to support the development of methods for genomic conservation.

## Author Contributions

RG, GAH, NRP, SO, and RJO designed the research. RG, NRP, and PJ performed DNA extractions, quality control, and sequencing. GAH and RG performed read analysis and classification. GAH and RJO performed mitochondrial assembly and phylogenetics. NMT performed methylation analysis. GAH, RG, NMT, and RJO wrote the manuscript and all authors contributed to manuscript revisions.

## Supplementary Figures

**Supplementary Figure 1:** The molecular weight of *X. caerulea* DNA, as measured by the Agilent Femto Pulse system, obtained from head and thorax extractions using one of three extraction protocols: the Qiagen DNeasy Blood & Tissue Kit, the Monarch HMW DNA Extraction Kit, and a modified phenol-chloroform extraction. DNA extractions are labeled according to the corresponding run code (Supplementary Table S2).

## Tables

**Supplementary Table 1.** A side-by-side comparison of the modified phenol-chloroform extraction method and the original protocol by Green and Sambrook (Fourth Edition).

**Supplementary Table 2.** Summary of extraction methods used for each run, including the number of specimens processed, the type and weight of tissue used, and the corresponding dsDNA concentrations measured by the Qubit 4 fluorometer. The table also provides the 260/280 and 260/230 DNA purity ratios for each run obtained from the Nanodrop One Spectrophotometer, and top DNA fragment length peaks from the Agilent FemtoPulse System (see Supplementary Figure 1). The italicized extractions (R5 and R6) were pooled for sequencing.

**Supplementary Table 3.** ONT sequencing run statistics. Run statistics for the *X. caerulea* PromethION run are listed. Corresponding statistics for reads identified to be endogenous DNA via alignment and BLAST classification are also shown.

**Supplementary Table 4.** Mapping statistics for *X. caerulea* data to public chromosome-level, *Xylocopa* assemblies. The total number of reads, number of unmapped reads, and number of primary aligned reads are shown.

**Supplementary Table 5.** Simplified BLAST classification groups of sequencing reads per a BLAST search against the full NCBI non-redundant nucleotide database. Reads matching the BLAST classification “bees,” “ants,” “wasps, ants, and bees,” and “hymenopterans” were considered a conservative estimate to identify endogenous, *Xylocopa*-derived reads.

**Supplementary Table 6.** Full BLAST classification groups of sequencing reads per a BLAST search against the full NCBI non-redundant nucleotide database. Reads matching the BLAST classification “bees,” “ants,” “wasps, ants, and bees,” and “hymenopterans” were considered a conservative estimate to identify endogenous, *Xylocopa*-derived reads.

**Supplementary Table 7.** *COX1* sequences obtained from NCBI and used to construct phylogenetic trees.

**Supplementary Table 8.** Global 5mC and 5hmC methylation and read depth characteristics across annotated genes.

**Supplementary Table 9.** High, mid, and low methylation patterns for select representative loci 2KB up and downstream of a gene. The table shows the coordinates used, depth, and percent for 5mC and 5hmC marks.

## Notes

### Competing Interest Statement

Rachel O'Neill serves on the Scientific Advisory Board of Colossal Biosciences.

## References

1. Allio, Rémi, Alex Schomaker-Bastos, Jonathan Romiguier, Francisco Prosdocimi, Benoit Nabholz, and Frédéric Delsuc. 2020. “MitoFinder: Efficient Automated Large-Scale Extraction of Mitogenomic Data in Target Enrichment Phylogenomics.” Molecular Ecology Resources 20 (4): 892–905.

2. Altschul, Stephen F., Warren Gish, Webb Miller, Eugene W. Myers, and David J. Lipman. 1990. “Basic Local Alignment Search Tool.” Journal of Molecular Biology 215 (3): 403–410.

3. Amarasinghe, Shanika L., Shian Su, Xueyi Dong, Luke Zappia, Matthew E. Ritchie, and Quentin Gouil. 2020. “Opportunities and Challenges in Long-Read Sequencing Data Analysis.” Genome Biology 21 (1): 30.

4. “Argk1 Arginine Kinase 1 [Drosophila Melanogaster (fruit Fly)] - Gene - NCBI.” n.d. Accessed April 3, 2026. https://www.ncbi.nlm.nih.gov/gene/39041.

5. “BuGZ Bub3 Interacting GLEBS and Zinc Finger Domain Protein [Drosophila Melanogaster (fruit Fly)] - Gene - NCBI.” n.d. Accessed April 3, 2026. https://www.ncbi.nlm.nih.gov/gene/35004.

6. Card, Daren C., Beth Shapiro, Gonzalo Giribet, Craig Moritz, and Scott V. Edwards. 2021. “Museum Genomics.” Annual Review of Genetics 55 (1): 633–659.

7. Cheetham, Seth W., Michaela Kindlova, and Adam D. Ewing. 2022. “Methylartist: Tools for Visualizing Modified Bases from Nanopore Sequence Data.” Bioinformatics (Oxford, England) 38 (11): 3109–3112.

8. De Coster, Wouter, and Rosa Rademakers. 2023. “NanoPack2: Population-Scale Evaluation of Long-Read Sequencing Data.” Bioinformatics (Oxford, England) 39 (5): btad311.

9. Deguines, Nicolas, Clémentine Jono, Mathilde Baude, Mickaël Henry, Romain Julliard, and Colin Fontaine. 2014. “Large-scale Trade-off between Agricultural Intensification and Crop Pollination Services.” Frontiers in Ecology and the Environment 12 (4): 212–217.

10. Dorado: Oxford Nanopore’s Basecaller. n.d. Github. Accessed April 2, 2026. https://github.com/nanoporetech/dorado.

11. Eeraerts, Maxime, Ruben Vanderhaegen, Guy Smagghe, and Ivan Meeus. 2020. “Pollination Efficiency and Foraging Behaviour of Honey Bees and Non-*Apis* Bees to Sweet Cherry.” Agricultural and Forest Entomology 22 (1): 75–82.

12. “Gek Genghis Khan [Drosophila Melanogaster (fruit Fly)] - Gene - NCBI.” n.d. Accessed April 3, 2026. https://www.ncbi.nlm.nih.gov/gene/37858.

13. Gerling, D., H. H. W. Velthuis, and A. Hefetz. 1989. “Bionomics of the Large Carpenter Bees of the Genus Xylocopa.” Annual Review of Entomology 34 (1): 163–190.

14. Green, Michael Richard, and Joseph Sambrook. 2012. Molecular Cloning: A Laboratory Manual. Cold Spring Harbor Laboratory Press.

15. Gu, Yifan, Wensu Han, Yuquan Wang, et al. 2023. “Xylocopa Caerulea and Xylocopa Auripennis Harbor a Homologous Gut Microbiome Related to that of Eusocial Bees.” Frontiers in Microbiology 14 (May): 1124964.

16. Gu, Zuguang, Lei Gu, Roland Eils, Matthias Schlesner, and Benedikt Brors. 2014. “Circlize Implements and Enhances Circular Visualization in R.” Bioinformatics (Oxford, England) 30 (19): 2811–2812.

17. Holmquist, Anna, Holly Tavris, Grace Kim, Lauren Esposito, Brian Fisher, and Athena Lam. 2025. “Towards Large-Scale Museomics Projects: A Cost-Effective and High-Throughput Extraction Method for Obtaining Historical DNA from Museum Insect Specimens.” Molecular Ecology Resources 25 (8): e14117.

18. Jeong, Hyeonsoo, Xin Wu, Brandon Smith, and Soojin V. Yi. 2018. “Genomic Landscape of Methylation Islands in Hymenopteran Insects.” Genome Biology and Evolution 10 (10): 2766–2776.

19. Junqueira, C. N., K. Hogendoorn, and S. C. Augusto. 2012. “The Use of Trap-Nests to Manage Carpenter Bees (Hymenoptera: Apidae: Xylocopini), Pollinators of Passion Fruit (Passifloraceae:*Passiflora Edulis*f.*flavicarpa*).” Annals of the Entomological Society of America 105 (6): 884–889.

20. Kearse, Matthew, Richard Moir, Amy Wilson, et al. 2012. “Geneious Basic: An Integrated and Extendable Desktop Software Platform for the Organization and Analysis of Sequence Data.” Bioinformatics (Oxford, England) 28 (12): 1647–1649.

21. LaBerge, Wallace E. 1965. “A Classification of the Large Carpenter Bees (Xylocopini) (Hymenoptera: Apoidea).Paul D. Hurd, Jr., J. S. Moure.” The Quarterly Review of Biology 40 (1): 97–97.

22. Lalueza-Fox, Carles. 2022. “Museomics.” Current Biology 32 (21): R1214–R1215.

23. Lewis, Samuel H., Laura Ross, Stevie A. Bain, et al. 2020. “ Widespread Conservation and Lineage-Specific Diversification of Genome-Wide DNA Methylation Patterns across Arthropods.” PLoS Genetics 16 (6): e1008864.

24. Leys, R., S. J. B. Cooper, and M. P. Schwarz. 2002. “Molecular Phylogeny and Historical Biogeography of the Large Carpenter Bees, Genus Xylocopa (Hymenoptera: Apidae).” Biological Journal of the Linnean Society. Linnean Society of London 77 (2): 249–266.

25. Li, Heng, Bob Handsaker, Alec Wysoker, et al. 2009. “The Sequence Alignment/Map Format and SAMtools.” Bioinformatics (Oxford, England) 25 (16): 2078–2079.

26. Lyko, Frank, Sylvain Foret, Robert Kucharski, Stephan Wolf, Cassandra Falckenhayn, and Ryszard Maleszka. 2010. “The Honey Bee Epigenomes: Differential Methylation of Brain DNA in Queens and Workers.” PLoS Biology 8 (11): e1000506.

27. Mensah, Ben, and Andreas Kudom. 2011. “Foraging Dynamics and Pollination Efficiency of *Apis Mellifera* and *Xylocopa Olivacea* on *Luffa Aegyptiaca* Mill (Cucurbitaceae) in Southern Ghana.” Journal of Pollination Ecology 4 (May): 34–38.

28. Miller, O. J., W. Schnedl, J. Allen, and B. F. Erlanger. 1974. “5-Methylcytosine Localised in Mammalian Constitutive Heterochromatin.” Nature 251 (5476): 636–637.

29. Modkit: A Bioinformatics Tool for Working with Modified Bases. n.d. Github. Accessed April 2, 2026. https://github.com/nanoporetech/modkit.

30. Nachman, Michael W., Elizabeth J. Beckman, Rauri Ck Bowie, et al. 2023. “Specimen Collection Is Essential for Modern Science.” PLoS Biology 21 (11): e3002318.

31. Putiri, Emily L., and Keith D. Robertson. 2011. “Epigenetic Mechanisms and Genome Stability.” Clinical Epigenetics 2 (2): 299–314.

32. Quinlan, Aaron R., and Ira M. Hall. 2010. “BEDTools: A Flexible Suite of Utilities for Comparing Genomic Features.” Bioinformatics (Oxford, England) 26 (6): 841–842.

33. Rahman, Sarthok Rasique, and Jeffrey D. Lozier. 2023. “Genome-Wide DNA Methylation Patterns in Bumble Bee (Bombus Vosnesenskii) Populations from Spatial-Environmental Range Extremes.” Scientific Reports 13 (1): 14901.

34. Raxworthy, Christopher J., and Brian Tilston Smith. 2021. “Mining Museums for Historical DNA: Advances and Challenges in Museomics.” Trends in Ecology & Evolution 36 (11): 1049–1060.

35. Rey, Olivier, Christophe Eizaguirre, Bernard Angers, et al. 2020. “Linking Epigenetics and Biological Conservation: Towards a Conservation Epigenetics Perspective.” Functional Ecology 34 (2): 414–427.

36. Seebacher, Frank, and Jens Krause. 2019. “Epigenetics of Social Behaviour.” Trends in Ecology & Evolution 34 (9): 818–830.

37. Shen, Wei, Botond Sipos, and Liuyang Zhao. 2024. “SeqKit2: A Swiss Army Knife for Sequence and Alignment Processing.” iMeta 3 (3): e191.

38. Stamatakis, Alexandros. 2014. “RAxML Version 8: A Tool for Phylogenetic Analysis and Post-Analysis of Large Phylogenies.” Bioinformatics (Oxford, England) 30 (9): 1312–1313.

39. Stavenga, Doekele G. 2023. “Pigmentary Colouration of Hairy Carpenter Bees, Genus Xylocopa.” The Science of Nature 110 (3): 22.

40. Uliano-Silva, Marcela, João Gabriel R. N. Ferreira, Ksenia Krasheninnikova, et al. 2023. “MitoHiFi: A Python Pipeline for Mitochondrial Genome Assembly from PacBio High Fidelity Reads.” BMC Bioinformatics 24 (1): 288.

41. Wang, Hongfang, Zhenguo Liu, Ying Wang, Lanting Ma, Weixing Zhang, and Baohua Xu. 2020. “Genome-Wide Differential DNA Methylation in Reproductive, Morphological, and Visual System Differences between Queen Bee and Worker Bee (Apis Mellifera).” Frontiers in Genetics 11 (August): 770.

42. Wickham, Hadley. 2016. Ggplot2: Elegant Graphics for Data Analysis. 2nd ed. Use R! Springer International Publishing. PDF.

43. Winfree, Rachael, James R. Reilly, Ignasi Bartomeus, Daniel P. Cariveau, Neal M. Williams, and Jason Gibbs. 2018. “Species Turnover Promotes the Importance of Bee Diversity for Crop Pollination at Regional Scales.” Science (New York, N.Y.) 359 (6377): 791–793.

44. Yagound, Boris, Emily J. Remnant, Gabriele Buchmann, and Benjamin P. Oldroyd. 2020. “Intergenerational Transfer of DNA Methylation Marks in the Honey Bee.” Proceedings of the National Academy of Sciences of the United States of America 117 (51): 32519–32527.

45. Zhang, Dan, Jianfeng Jin, Zeqing Niu, et al. 2025. “Chromosome-Level Genome Assembly of the Large Carpenter Bee Xylocopa Dejeanii Lepeletier, 1841 (Hymenoptera: Apidae).” Scientific Data 12 (1): 1280.

46. Zimmermann, Juergen, Mehrdad Hajibabaei, David C. Blackburn, et al. 2008. “DNA Damage in Preserved Specimens and Tissue Samples: A Molecular Assessment.” Frontiers in Zoology 5 (1): 18.

