## Supplemental Figure 1 for "Optimization of High Molecular Weight DNA Extractions from Dried, Museum-Grade Insects Enables Long-Read Sequencing, Phylogenetics, and Methylation Profiling"

Hartley, Green et al Supp Figure 1

### Qiagen DNeasy Blood and Tissue Kit

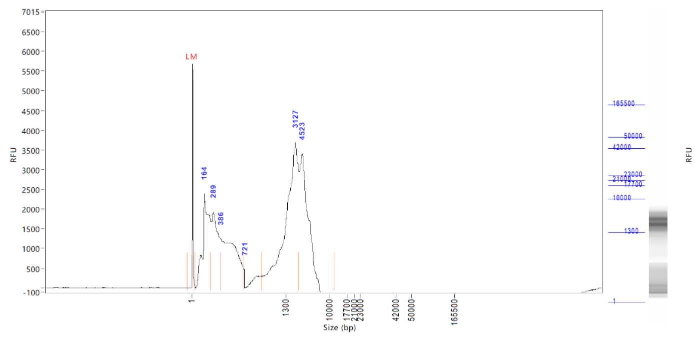


3x head (R1)

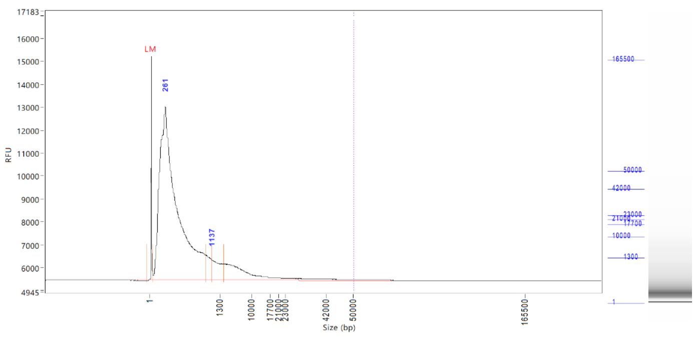


3x thorax (R2)

### Monarch HMW DNA Extraction Kit

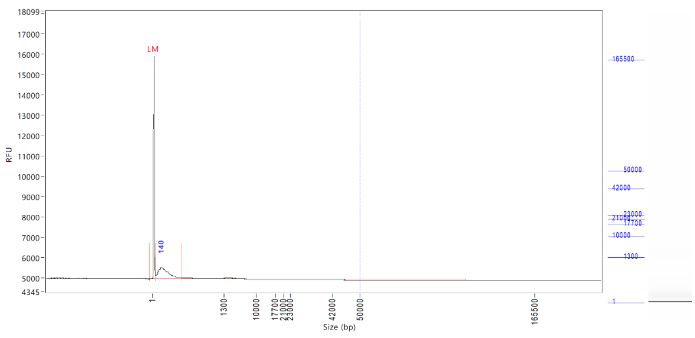


3x head (R3)

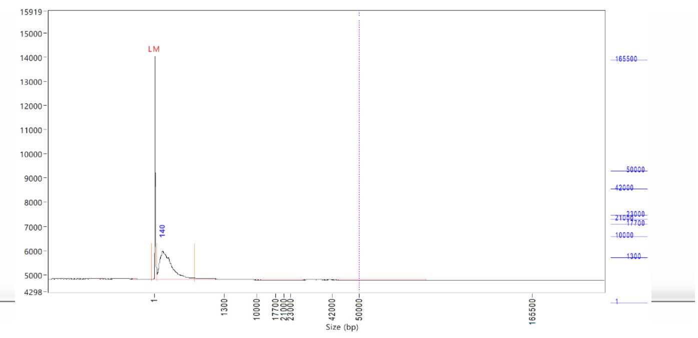


3x thorax (R4)

### Phenol-Chloroform Extraction

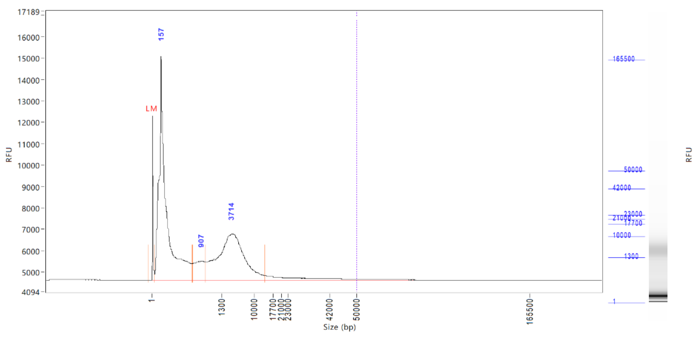


1x head (R5)

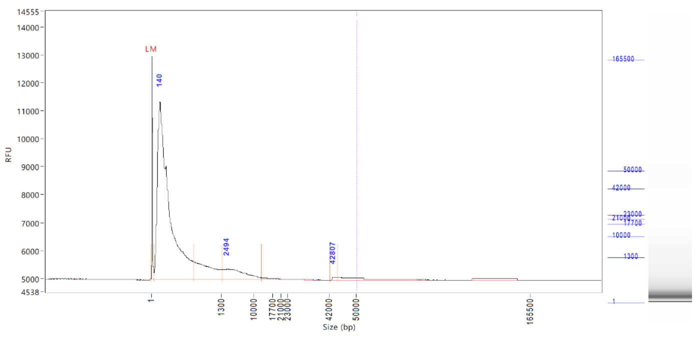


1x thorax (R6)

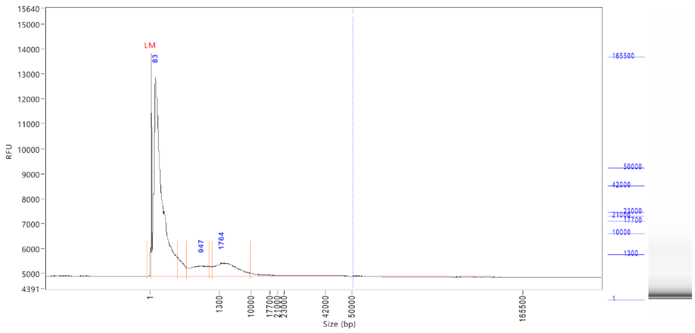


1x head (R8)

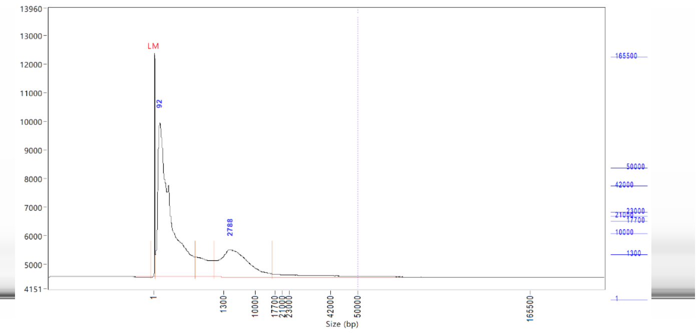


3x thorax (R9)
